# Rapid actin filament turnover maintains cortical connectivity while allowing for cell cortex deformation and flow

**DOI:** 10.64898/2026.05.24.727551

**Authors:** Rachel S. Kadzik, Ondrej Maxian, Isaiah Thomas, David R. Kovar, Edwin M. Munro

## Abstract

Cells harness the actomyosin contractility of the cell cortex to drive rapid cellular deformations and intracellular flows during cell polarization, migration, and division. To sustain contractile network architectures while allowing for network deformation and remodeling, the balance of actin filament assembly and disassembly must be finely tuned, but how this is coordinated in the cell remains obscure. Here, we combine quantitative measurements and manipulations of filament assembly and disassembly rates with live imaging of network contractility dynamics in the *C. elegans* zygote to identify co-dependencies between assembly rates, disassembly rates, and large-scale deformations of the cortical actin network. We find that strong reductions in either filament assembly or disassembly rates both result in actin cortex collapse, but each perturbation has distinct effects on actin cortex and cell membrane dynamics. These findings demonstrate that rapid turnover, involving tightly coordinated assembly and disassembly, allows the cortex to maintain a connected architecture while undergoing rapid deformation and coherent flow.

## Introduction

Just beneath the plasma membrane of the cell, a network of actin filaments, myosin mini-filaments, and actin binding proteins form the cell cortex that allows the cell to maintain and alter its shape (Chugh and Paluch, 2018). While the molecular properties of many of the individual components that constitute the actin cell cortex are well characterized (Lappalainen et al., 2022; Blanchoin et al., 2014), understanding how these players interact to affect large scale behaviors on the network level continues to be a challenge (Keren et al., 2008; Naganathan et al., 2018; Chugh et al., 2017).

An important function of the actomyosin cell cortex is generation of myosin-mediated cortical flows that contribute to polarization of the cell (Munro et al., 2004; Oon and Prehoda, 2019; Hird and White, 1993; Mayer et al., 2010; McFadden et al., 2017) and formation of the cytokinetic ring (White and Borisy, 1983; Reymann et al., 2016; Li and Munro, 2021; Khaliullin et al., 2018). Transmission of coherent cortical flows requires coordination of actin filament length, density, architecture, and crosslinking (Chugh et al., 2017; Ierushalmi et al., 2020), while dissipation of accumulated contractile strain is needed to maintain network integrity and prevent cortex rupture(Chandrasekaran et al., 2019; Hiraiwa and Salbreux, 2016; Mak et al., 2016; McFadden et al., 2017; Popov et al., 2016)). Untangling how these many factors interact to facilitate formation and maintenance of cortex dynamics can be daunting, and through a combination of *in vitro* (Murrell and Gardel, 2012; Stam et al., 2017; Freedman et al., 2019; Sakamoto et al., 2023), *in silico* (Freedman et al., 2017; Popov et al., 2016; Mak et al., 2016; Belmonte et al., 2017; McFadden et al., 2017; Jung et al., 2019; Yu et al., 2018), and *in vivo* (Fritzsche et al., 2013; Naganathan et al., 2018; Mayer et al., 2010; Chugh et al., 2017; Koenderink and Paluch, 2018; Salbreux et al., 2012; Ofer et al., 2011; Raz-Ben Aroush et al., 2017) investigations the field has gained insight into the requirements for this process. What has emerged from this work from many labs is the importance of the turnover of actin filaments and network components to allow for both cortex remodeling and release of buildup of contractile strain (Chandrasekaran et al., 2019; Hiraiwa and Salbreux, 2016; Mak et al., 2016; McFadden et al., 2017; Popov et al., 2016) needed to facilitate cortical flow.

The cell cortex of the one-cell C. elegans embryo exhibits a variety of actin architectures and patterns of cortical flows that serve to polarize PAR components (Munro et al., 2004; Costache et al., 2022; Mayer et al., 2010; Hird and White, 1993) and complete cytokinesis (Reymann et al., 2016; Li and Munro, 2021). This system has been used to highlight proteins that have similar molecular signatures on large-scale morphological changes (Naganathan et al., 2018), but how specific actin interacting proteins may be coordinating or antagonizing their activities to tune cortex properties to enable long-range flow remains unclear. To gain insight into how turnover modulates actin cortical dynamics *in vivo*, we focused on the two major factors that dictate actin filament turnover dynamics: filament nucleation and assembly balanced against filament disassembly and actin monomer recycling. We specifically focused on how disrupting turnover either by slowing assembly or slowing disassembly affects cortex and membrane dynamics as well as feeds back on assembly and disassembly rates.

Making use of single molecule imaging and particle tracking pipelines, we developed sensitive methods for quantifying actin filament assembly and disassembly in vivo. Using graded depletion of either profilin, to slow linear actin assembly, or cofilin to slow filament disassembly, we have been able to evaluate the effects of modulating these aspects of filament turnover on actin cortex and cell membrane dynamics in vivo. Additionally, co-dependencies between filament assembly and disassembly emerged, providing additional insight into how the cell finely tunes turnover dynamics to maintain cortex integrity and cortical flow.

## Results

### *In vivo* quantification of formin-mediated filament assembly at the actomyosin cortex

To determine how modulating the inputs to filament turnover affects actomyosin cortical dynamics, we first needed to establish sensitive *in vivo* methodology to quantify rates of F-actin assembly and disassembly in the *C. elegans* cortex. We started by focusing on quantification of F-actin assembly rates. Although elongation of individual actin filaments cannot be visualized in animal cells, tagged proteins that associate with elongating filaments can be observed. Using an endogenously fluorescently labeled diaphanous-related formin, CYK-1::GFP, allowed us to quantify rates of formin-mediated filament assembly at the cortex (Padmanabhan et al., 2017; Li and Munro, 2021). Formins form dimers on the barbed end of elongating actin filaments and increase the rate of elongation in the presence of profilin-actin (schematized in Fig 1A) (Kovar et al., 2006; Neidt et al., 2009; Oosterheert et al., 2024). Loss of CYK-1::GFP dramatically effects cortical actin assembly (Swan et al., 1998; Severson et al., 2002), and we predict CYK-1 makes a substantial contribution to the cortical actin network in the *C. elegans* zygote.

**Figure 1.**
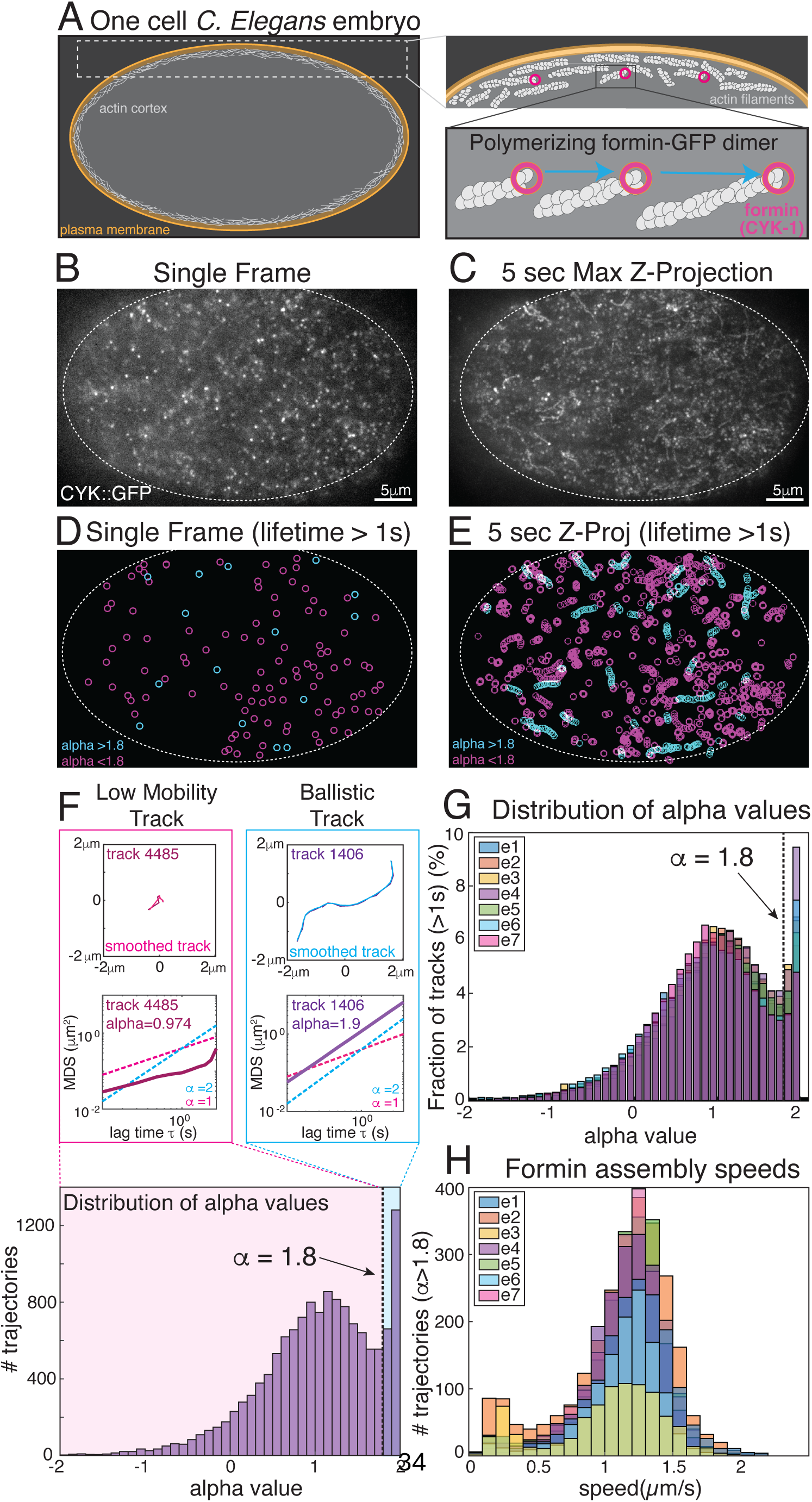
Particle tracking of tagged formin in actin cortex to quantify formin assembly speeds in the one cell *C. elegans* zygote. **(A)** Schematic of the actin cortex in the one cell embryo. Formin dimers polymerize filamentous actin in the cortical plane, which can be visualized by near-TIRF microscopy. **(B)** Representative image from TIRFM timelapse (Video 1) of endogenously tagged formin, CYK-1::GFP. **(C)** 5 second maximum z-projection from Video 1 showing CYK-1::GFP particle movement over a 5 second window. **(D)** Single frame (as in B) with all particles with lifetime over 1 second in length tracked. Magenta circles are low-mobility tracks and cyan circles are ballistic tracks. **(E)** 5 second maximum z-projection (as in C), magenta circles are low mobility tracks and cyan circles are ballistic tracks. **(F)** Representative low mobility or ballistic track from embryo above. Low mobility tracks (magenta, track 4485) are characterized by small amount of displacement over time, and an anomalous diffusion exponent (a) close to 1. Ballistic tracks (cyan, track 1406) are characterized by large displacements over time, directed motion, and an anomalous diffusion exponent closer to 2. Distribution of a-values for all particles tracked (15949) for longer than 1 second display two clear peaks, one broad peak at an alpha value close to 1 and a sharp peak near 2. We set the cutoff for ballistic motion at an alpha value of 1.8 to restrict our analysis to particles that only exhibit directed, progressive motion. **(G)** Distribution of alpha values for 7 control embryos, 122,715 total tracks analyzed. **(H)** Distribution of formin particle speeds only for tracks exhibiting ballistic motion (alpha values greater or equal to 1.8, 12,161 tracks, 7 embryos), with a mean formin speed at the cortex of 1.13 +/- 0.33 mm/sec.

Visualizing CYK-1::GFP at the *C. elegans* cortex using near-TIRF microscopy shows numerous diffraction-limited particles, likely representing formin dimers (Fig 1B and Video 1). Each CYK-1::GFP particle can be detected using a combined filter for fluorescent intensity and size and tracked over time (Fig 1B). The particles trajectories show a range of mobility patterns, from particles that demonstrate very little movement over a 5 second window (low mobility particles, magenta, Fig 1D, E, F) to particles that have persistent, directed motion (ballistic particles, cyan, Fig 1D, E, F). We tracked all detected particles with lifetimes over 1 second (20 frames) and filtered the resulting tracks based on the anomalous diffusion coefficient, alpha, evaluating the mean squared displacement versus time lag tau. Tracks with alpha values close to 1 display characteristic diffusive motion, and tracks with alpha values approaching 2 exhibit ballistic motion. Representative tracked particles of low-mobility and ballistic track trajectories (Fig 1F) and alpha values from the pictured embryo (Fig 1B, C) show characteristic tracks for each type. Alpha values between 1 and 2 can have a range of trajectory patterns: from movement due to cortical flow, to trajectories with both ballistic and static portions, to trajectories exhibiting only ballistic motion. To minimize effects of cortical flow and tearing on our analysis, we restricted our analysis to the mitotic phase of the cell cycle, which exhibits relatively slow cortical flow and minimal cortical tearing. Plotting the alpha values of all the trajectories shows two prominent peaks, one broad peak at 1, representing low mobility CYK-1::GFP at the cortex, and a second peak at approximately 1.8-2. This distribution of alpha values is highly reproducible across multiple embryos, as seen in Fig 1G (7 embryos, 122,715 tracks).

To restrict our analysis to trajectories consisting of primarily ballistic motion, we set a threshold for alpha value of greater or equal to 1.8. (Fig 1G). Even with this strict threshold value for alpha, we still have hundreds of tracks per embryo for further evaluation. Using minimal smoothing to determine the mean squared displacement over time, we calculate the trajectory speed for those trajectories with alpha values greater than 1.8. This results in a distribution of elongation speeds (Fig 1H) which is highly reproducible between multiple embryos, with a mean elongation speed of 1.13 +/-0.33 μm/sec (standard deviation). These speeds are very similar to previously reported CYK-1 elongation speeds in the *C. elegans* embryo (1.1-1.3 μm/sec), using similar methodology for trajectory speed calculation, but an independently generated endogenous CYK-1::GFP strain (Costache et al., 2022). These values are somewhat slower than the range of formin speeds (1.98- 4.9 μm/sec) reported in mammalian cells (Funk et al., 2019).

### *In vivo* quantification of filament disassembly at the actomyosin cortex

With highly reproducible methods for quantifying formin elongation rates in the zygote cortex we moved to adapting methods to evaluate actin filament disassembly *in vivo* to get a complete picture of actin filament turnover in the cell. We modified a previously used method for tracking single actin particles *in vivo* to determine actin lifetime at the cortex (Robin et al., 2014; Li and Munro, 2021; Michaux et al., 2018). Using a transgenic actin::GFP strain (Robin et al., 2014) that expresses low levels of actin::GFP, we used near-TIRF microscopy to visualize actin::GFP at the cortex at single molecule levels (Fig 2B). Actin::GFP molecules that are at the cortex for at least 2 seconds (2 frames) are presumed to incorporate into actin filaments, and actin::GFP with lifetimes less than 2 seconds are assumed to be actin monomers only transiently present at the cortex (Fig 2A). Using single molecule detection, individual actin::GFP particles were identified and tracked over time (Fig 2B and Video 2). Kymographs of selected particles shows the range of lifetimes at the cortex for selected particles (Fig 2C). Tracking all the actin::GFP particles in an embryo during mitosis and plotting the change in number of particles at the cortex over time gives a survival curve demonstrating how the number of actin::GFP molecules at the cortex changes over time (Fig 2D). These curves are highly reproducible and show the disappearance rate of actin::GFP at the cortex. Consistent with previous work, we assumed that actin::GFP disappearance from the cortex is due to a combination of photobleaching of the fluorophore and F-actin disassembly (Watanabe et al., 2018; Robin et al., 2014). By collecting data at different duty ratios, we can calculate the fluorophore photobleaching rate and the actin::GFP off rate (see Materials and Methods, (Robin et al., 2014)).

**Figure 2.**
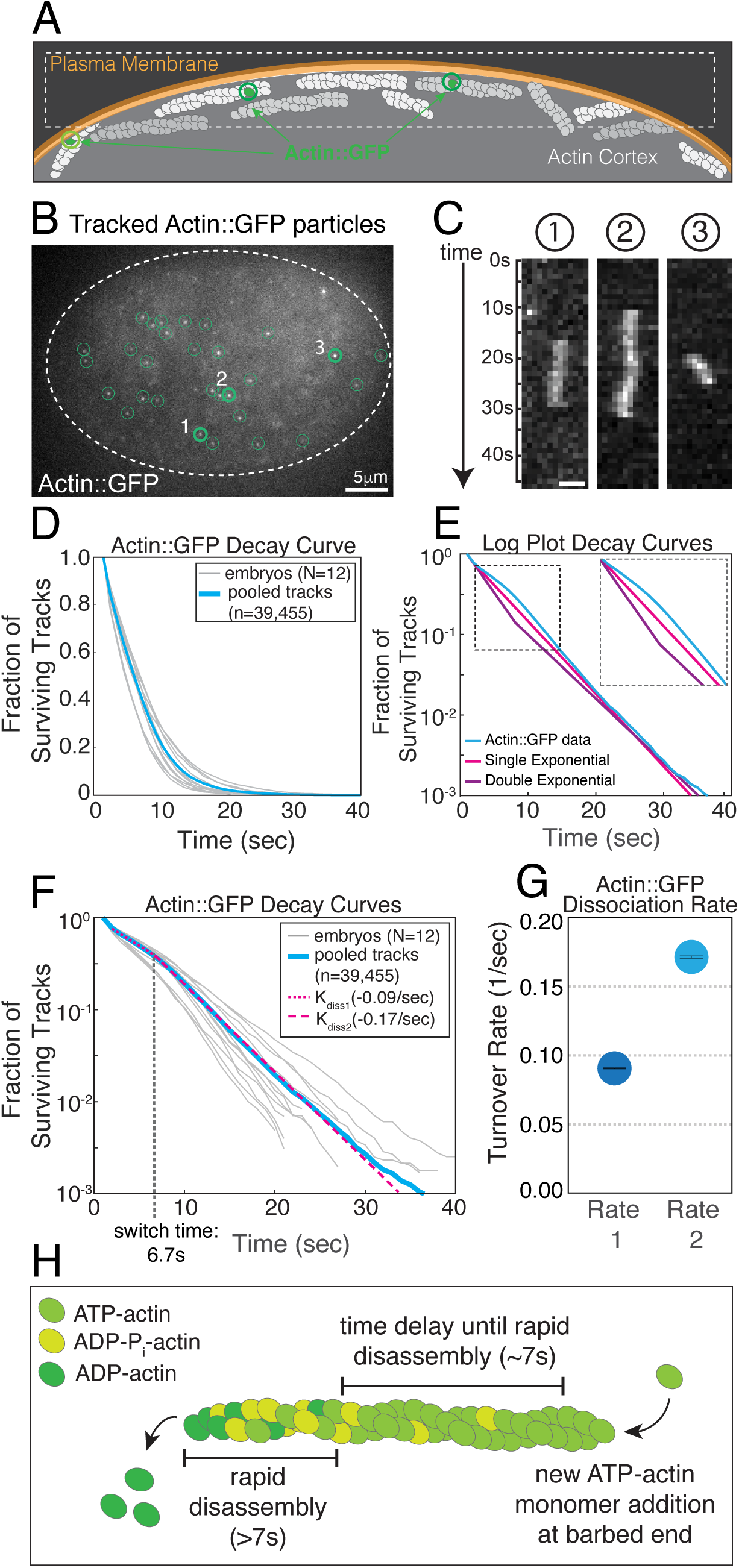
Particle tracking of actin::GFP molecules in the actin cortex to quantify F-actin turnover rates. **(A)** Schematic of actin cortex with sparse labeling of actin:GFP incorporated into actin filaments. **(B)** Representative image from TIRFM timelapse (Video 2) of sparsely labeled actin::GFP at the cortex, all tracked particles circled, representative particles 1, 2 and 3. **(C)** Kymographs of representative individual actin::GFP particles from B showing lifetime at the cortex. **(D)** Release curves for Actin::GFP molecules imaged with fixed laser power at low duty ratio (1 second interval). Thin gray traces show release curves for each embryo (N=12). Thick blue curve show data pooled for all embryos (n=39,455 tracks). **(E)** Actin::GFP release curve of pooled data (blue) plotted on a log10 plot to visualize the fit of exponential curves to the data. Representative single exponential and double exponential curves are plotted as reference. Zoom of the first 15 seconds in inset shows that neither a single exponential (pink) nor a double exponential (purple) fit the actin::GFP release curves in their entirety. **(F)** Dashed lines indicate the two independent single exponential fits to either the 1^st^ or 2^nd^ portion of the curve with a characteristic switching time. The two dissociation rates are corrected for photobleaching based on high laser duty ratio imaging (100ms, continuous acquisition) to determine the photobleaching rate. **(G)** Initial slow actin::GFP dissociation rate and secondary fast actin::GFP dissociation rate plotted with 95% confidence intervals obtained from bootstrap sampling. **(H)** Schematic of actin filament assembly, aging, and disassembly that would fit the two observed actin::GFP dissociation rates.

Although the decay curves appear to display mono-exponential decay kinetics, as would be expected if there were a single dissociation rate, we found that a single exponential (Fig 2E, magenta line) did not fit the observed data (Fig 2E, cyan line), suggesting that there is not a single off-rate for actin::GFP at the cortex. Alternatively, another common model for particle dissociation would be if the actin::GFP was a mixed population, with two different dissociation rates. In this case, we would expect a double exponential to fit the data (blue line in Fig 2E) where an initial fast rate predominates early, and slower off rate predominates later. Instead, the data fits an initial slow off rate, with a characteristic switching time of approximately 7 seconds, then a faster off rate after 7 seconds has elapsed (Fig 2F, 2G). The predominance of an initial slow rate of dissociation suggests that there is a block to the fast rate of dissociation for approximately 7 seconds of actin::GFP lifetime at the cortex, after which the actin filament is presumably rapidly disassembled.

This result was initially surprising as when we had previously performed analysis of actin::GFP dynamics at the cortex we had thought a single exponential fit matched the data reasonably well (Robin et al., 2014; Li and Munro, 2021). Furthermore, it might reasonably be expected that the actin::GFP represents a mixed population with different disassembly rates (Li and Munro, 2021; Pollard and Borisy, 2003; Fritzsche et al., 2013, 2016) which might fit double exponential kinetics, as the actin cortex is made up of linear, bundled and branched F-actin that is bound by different actin binding proteins that can modulate dynamics (Chugh et al., 2017; Blanchoin et al., 2014). But when considering the dissociation dynamics that best fit our observed actin::GFP behavior, we realized these results fit with a long-standing model for actin disassembly (Carlier, 1990; Theriot and Mitchison, 1992; Pollard and Borisy, 2003) (Fig 2H). Specifically, ATP-actin monomers are added to the growing barbed end of an actin filament, and it has long been appreciated *in vitro* that there is an initial block to rapid filament disassembly due to the nucleotide state of actin-ATP. Both the cleavage of ATP into ADP-Pi and the slower release of Pi need to occur to allow for a structural change in the filament that promotes filament disassembly (Pollard, 1986; Carlier and Pantaloni, 1986; Carlier, 1987; Jégou et al., 2011; Blanchoin and Pollard, 2002). While we cannot observe the multistep ATP hydrolysis and subsequent Pi release, the initial block to rapid disassembly that we observe suggests that we can quantify this phenomenon *in vivo*. *In vitro*, the slow step of Pi release has been measured at ∼300s, much longer than what we observe *in vivo*. It has long been noted that due to the observed rate of turnover of actin networks in cells this process of Pi release must occur faster *in vivo*, possibly through the activities of additional actin binding proteins such as cofilin or coronin (Goode et al., 2023).

### Small filament bundle dynamics integrate filament assembly and disassembly dynamics observed in single-particle analysis

While particle tracking in the cell cortex of either formin or actin particles allows for sensitive quantification of either formin-mediated F-actin assembly or all actin filament disassembly, it does not provide an integrated view of assembly and disassembly on the filament level. While we cannot visualize turnover dynamics on the single filament level, we found that we could visualize these dynamics in small F-actin bundles.

Plastin (PLST-1) is an actin binding protein that bundles actin filaments in the C. elegans actomyosin cortex (Ding et al., 2017; White et al., 2025). Imaging endogenously tagged PLST-1::GFP via TIRFM reveals that much of the actin cortex in the one-cell embryo is bundled by plastin (Fig 3A and Video 3), as observed by other groups (Ding et al., 2017; White et al., 2025) (Video 4). In the PLST-1::GFP cortex, we consistently noted plastin ‘streaks’ in the cortex characterized by the initial appearance of an elongating plastin bundle, a perdurance of the plastin signal for a set length of time, then a disappearance of the bundle from the non-polymerizing end (Fig 3B). By tracing these streaks in the cortex and generating a kymograph, we can visualize the growing end of the plastin bundle, and after a set period of time, a progressive disappearance of the bundle from the non-elongating end. These dynamics are reminiscent of filament treadmilling with a bundle assembly rate closely tied to bundle disassembly rate after a set bundle lifetime (Fig 3C).

**Figure 3.**
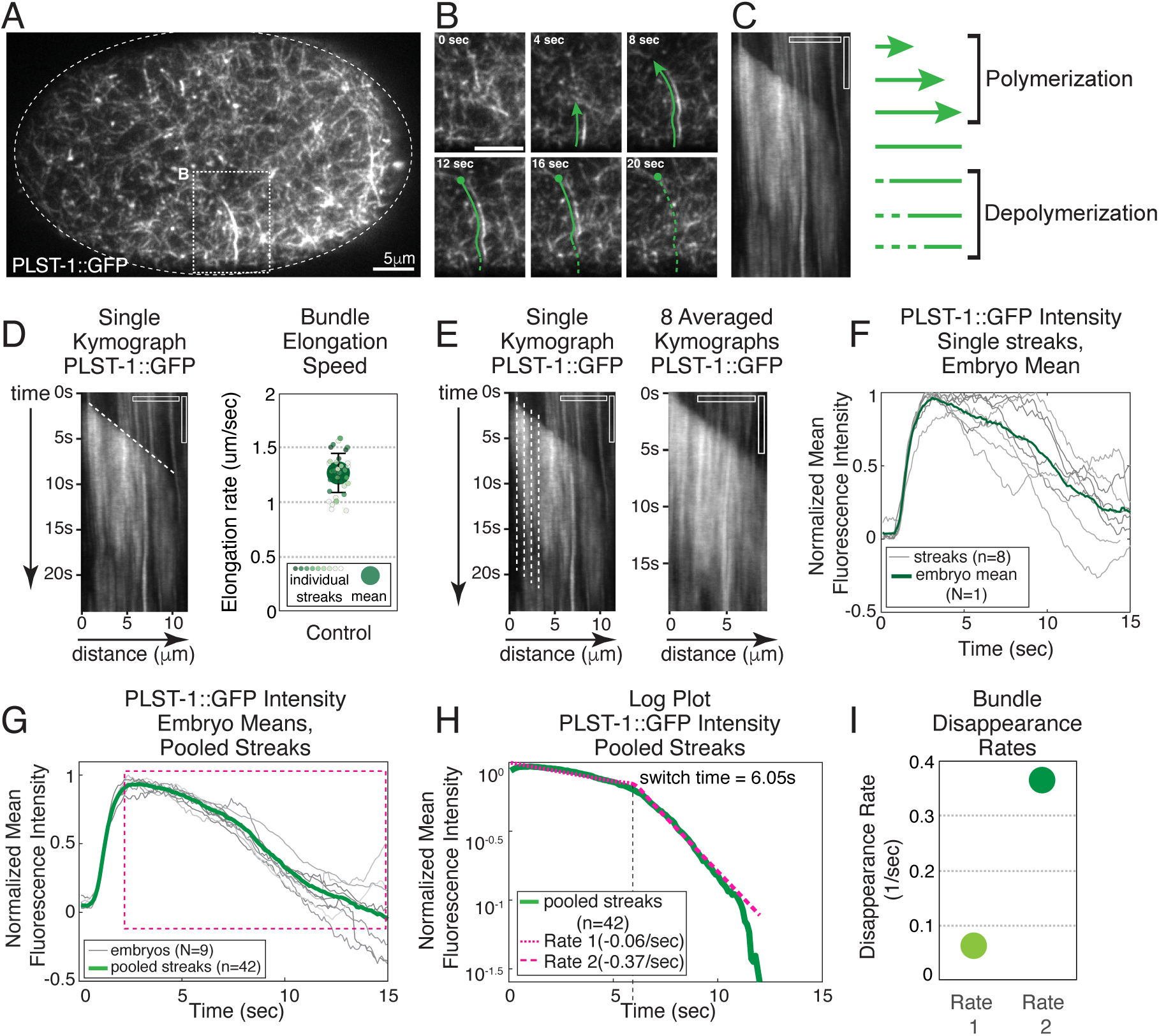
Plastin-GFP filamentous actin bundle dynamics in the actin cortex. **(A)** Representative image from TIRFM timelapse (Video 3) of endogenously tagged plastin (PLST-1::GFP) in the cortex of the one cell *C. elegans* zygote. **(B)** Time series of plastin streak in the actin cortex. **(C)** Kymograph of streak in B, with time on the y axis and distance on the x axis. Plastin appearance over time represents bundle elongation rate, and subsequent disappearance after a set period of time. **(D)** Plastin bundle elongation rate can be determined by measuring the slope of the PLST-1::GFP appearance over time. The mean elongation rate is 1.27 +/- 0.18 mm/sec, close to the formin assembly rates measured from CYK-1::GFP tracked particles (Fig 1). N=9 embryos, n= 42 streaks **(E)** Quantification of plastin lifetime and disassembly is averaged over the entire kymograph, lining up the start at each point along the bundle at first appearance of plastin signal. 8 kymographs of plastin bundles from the embryo pictured in A were lined up and the averaged z-projection is displayed. **(F)** Normalized mean fluorescence intensity over time plotted for each plastin bundle (thin gray lines, n=8) and the embryo mean (thick green line) for embryo pictured above. **(G)** Plot of normalized mean fluorescence intensity over time plotted for each embryo (thin gray lines, N=9, each embryo had at least 2 plastin bundles) and all streaks from all embryos pooled (thick green line, n=42). **(H)** Log plot of normalized mean fluorescence intensity over time for inset in G. Similar to what is observed with the actin::GFP data, there is an initial slow disappearance rate, then a rapid disappearance rate after 6 minutes. **(I)** Plot of initial and secondary disappearance rates for plastin bundles at the cortex. All scale bars are 5mm for distance and 5 seconds for time (on kymographs).

To characterize the turnover dynamics of these plastin streaks, we measured both the assembly rate and disassembly dynamics. By measuring the slope of the plastin streaks from the kymographs, we determined a bundle polymerization rate of 1.27+/- 0.18 μm/sec (Fig 3D), strikingly similar to the CYK-1 polymerization rates quantified in Fig 1H (1.13 +/-0.33 μm/sec). Using 2-color imaging, we can visualize CYK-1::Halo at the tips of plastin streaks, confirming these streaks are generated through formin polymerization (Supplemental Fig 1, Video 5). To characterize the disassembly dynamics of the plastin streaks, we generated kymographs for each streak, then plotted the mean fluorescence intensity of the signal over time, lining up each column with the initial rise in fluorescence intensity corresponding with the growing bundle end. This allowed us to average the dynamics of the plastin signal (appearance and disappearance) at every point along the streak to generate a mean fluorescent intensity profile for each streak (Fig 3E, F). We can generate an averaged kymograph of 8 individual streaks from the same embryo (Fig 3F, green line), plot each streak (Fig 3F, gray lines) and the average streak intensity profile for one embryo (Fig 3F, green line). We can then plot individual profiles for multiple embryos (Fig 3G, gray lines) and the average for all embryos (N=9, Fig 3G, green line). While there is significant variability from streak to streak, averaging the streaks in an embryo provide a clear picture of the dynamics of the plastin signal over time, with characteristic rapid appearance, a reproducible time at the cortex, then rapid disappearance. Plotting the portion of the curves immediately following cessation of elongation on a log plot (Fig 3H), similar to the plot used to characterize the actin::GFP turnover dynamics, we see a similar shaped plot to the actin::GFP decay curves with an initial slow rate of decay, then a more rapid rate of decay after a set period of time (∼6 sec). While the rates of signal loss at the cortex are different between the single molecule actin::GFP and the PLST-1::GFP data (Fig 3I), the overall pattern of initial slow rate of disappearance followed by a more rapid rate of disappearance after a set period of time (approximately 6-7 seconds) is strikingly similar (Fig 3H). The PLST-1::GFP streak data provide a more complete picture of actin dynamics at the cortex and reinforce the findings from our single molecule analysis of cortical actin dynamics quantifying actin turnover. Plastin streaks provide an integrated view of actin bundle assembly and disassembly at the cortex and clearly demonstrate the close association of filament assembly and disassembly dynamics.

### Decreasing formin mediated F-actin assembly, or cofilin-mediated disassembly, slows filament turnover dynamics

Having established quantitative methods for evaluating actin filament turnover *in vivo*, we next wanted to investigate how reducing either filament assembly rates or filament disassembly rates affected cortical actin dynamics (Fig 4A). To specifically slow formin-mediated filament assembly, we focused on reducing expression of the predominant profilin in the early embryo (PFN-1) (Polet et al., 2006; Neidt et al., 2009) as diaphanous-related formins increase filament polymerization rates in the presence of profilin-actin. To reduce filament turnover, we knocked down the only identified cofilin isoform expressed in the early embryo, UNC-60A (Ono, 1999; Ono and Benian, 1998; Ono et al., 2003; Yamashiro et al., 2005). For either cofilin (UNC-60A) or profilin (PFN-1), severe knockdown of each component by feeding RNAi for over 26 hours caused cortical instabilities, cortex collapse and cytokinesis failure (Video 6). But, by feeding young adult worms with RNAi for different lengths of time (16-28 hours), we could generate a range of knockdown strengths in the developing embryos from mild, to moderate, to strong. We focused on evaluating these graded knockdown phenotypes to determine how a progressive loss of formin-mediated elongation (profilin/PFN-1) or filament disassembly (cofilin/UNC-60A) effected actin cortex dynamics. In evaluating these knockdowns, in addition to the timing, we could also score the knockdown based on overall phenotype, which could be appreciated even in single-molecule TIRF imaging. For both RNAi conditions, the mildest knockdowns did not have any obvious phenotypic differences from control. Moderate knockdowns had contractile instabilities and membrane blebbing at times of high myosin activity, such as in interphase and cytokinesis, and the strongest knockdowns exhibited increased instabilities and membrane blebbing throughout the cell cycle. We restricted our analysis of filament assembly and disassembly rates to mitosis, as in most knockdown conditions, there was little effect on overall cortex dynamics during this portion of the cell cycle. For embryos where the cortical network dynamics were too severely affected in mitosis, these embryos were left out of analysis, or we made use of NMY-2 RNAi to suppress myosin activity and maintain cortex integrity (Fig 4B).

**Figure 4.**
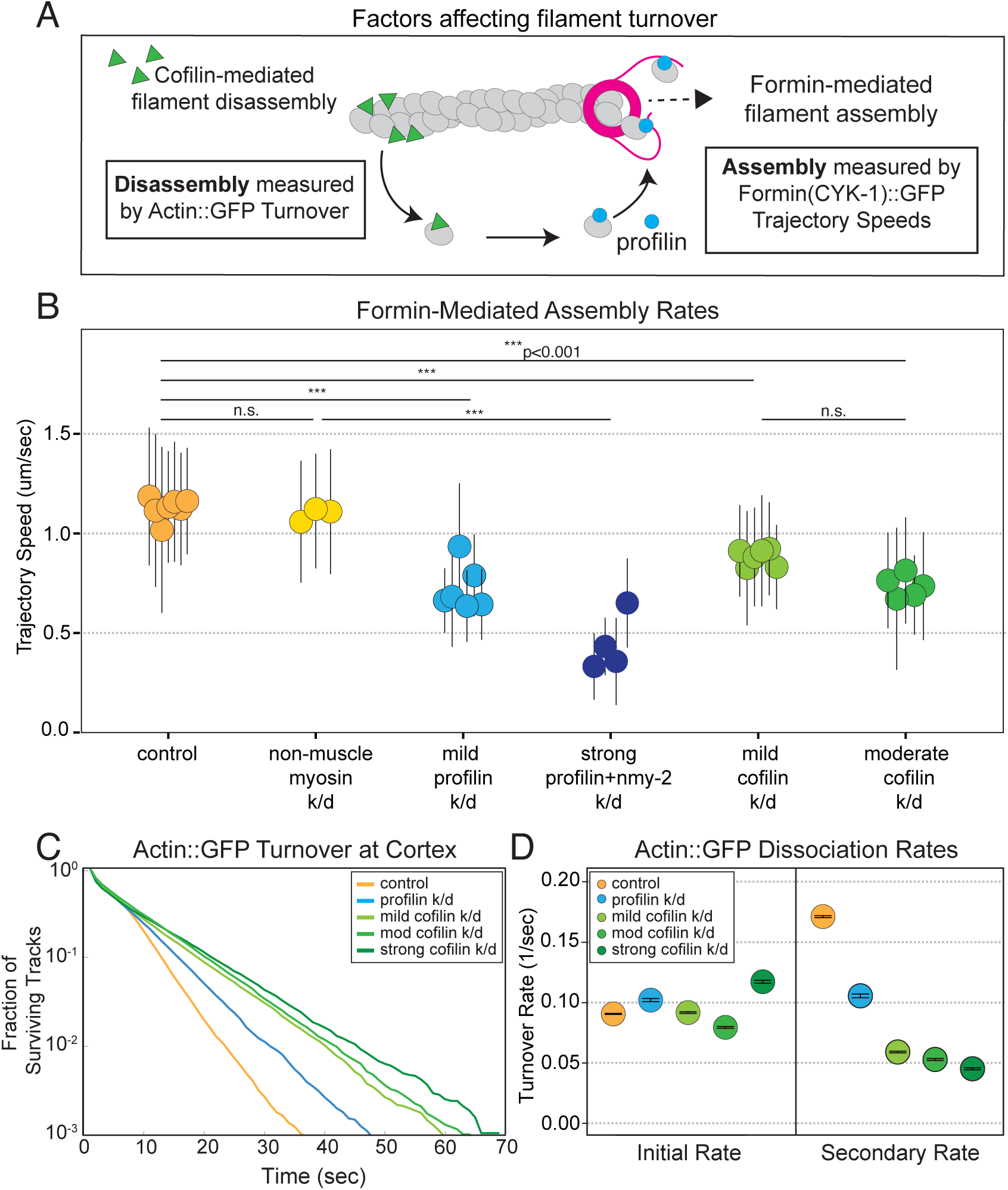
Graded knockdown of profilin or cofilin affect formin assembly rates and filamentous actin disassembly rates in the cortex. **(A)** Schematic of actin filament turnover, highlighting how either profilin or cofilin are expected to contribute to actin filament assembly and disassembly. **(B)** Measured trajectory speeds for RNAi knockdown of non-muscle myosin, profilin and cofilin. Each circle represents mean trajectory speed for 1 embryo, at least 500 trajectories analyzed per embryo. Error bars show standard deviation. Statistical significance was determined using pair-wise comparison using a mixed-effects model. **(C)** Log plot of pooled actin::GFP particle tracking release curves for control (orange, N=12), profilin knockdown (blue, N=6), mild cofilin knockdown (light green, N=5), moderate cofilin knockdown (green, N=3), and strong (dark green, N=2). **(D)** Initial slow actin::GFP dissociation rate and secondary fast actin::GFP dissociation rate calculated from two-slope fitting and photobleach correction for data in (C), means plotted with 95% confidence intervals using bootstrap sampling.

We started our analysis by looking at how formin-mediated assembly speeds were affected by graded knockdown of either profilin or cofilin. Highlighting the sensitivity of our single molecule analysis, even mild knockdown of profilin caused a statistically significant decrease in formin trajectory speeds (Fig 4B, light blue circles). Due to contractile instabilities with strong knockdown of profilin, non-muscle myosin knockdown was paired with strong profilin knockdown to allow for analysis of formin trajectories. NMY-2 knockdown alone does not have an appreciable effect on formin assembly speeds (Figure 4B, yellow circles). Strong knockdown of profilin paired with knockdown of NMY-2 resulted in a significant slowing of formin trajectory speeds, less than half the assembly rate observed in control or reduced NMY-2 conditions (Fig 4B, dark blue circles).

Somewhat unexpectedly, both mild and moderate knockdown of cofilin (Fig 4B, light and dark green circles) also reduced formin assembly speeds, although not to the degree of the profilin knockdown. We hypothesize that this decrease in formin trajectory speeds is due to decreased turnover of actin filaments, and the concomitant decrease in polymerization-competent actin monomers available for new filament assembly. While this result may be expected based on a limited availability of actin monomers in the cell, this is the first time that this tight relationship between assembly and disassembly kinetics has been demonstrated *in vivo*.

Upon finding a tight interconnected relationship between filament assembly and disassembly on formin-mediated F-actin assembly, we next wanted to evaluate the effect of decreasing either profilin levels or cofilin levels on F-actin disassembly rates. If our proposed model of the molecular mechanism underlying actin::GFP kinetics at the cortex (Fig 2H) is correct, we would predict cofilin (UNC-60A) to have a significant effect on the second, fast rate of actin::GFP dissociation, as cofilin preferentially binds to, and promotes severing and depolymerization at regions of ADP-actin (Blanchoin and Pollard, 1999; Suarez et al., 2011; Elam et al., 2013). With increased knockdown of cofilin, we observe a progressive slowing of the rate of dissociation of actin::GFP from the cortex, as predicted (Fig 4C and D, green lines and circles). Comparing the fast dissociation rates of actin::GPF, control exhibits of a predominant rate of 0.17/sec, mild cofilin 0.059/sec, moderate cofilin 0.053/sec and strong cofilin knockdown 0.045/sec. Of note, in the cofilin knockdown conditions, these slopes are generally uniform both early and late, not showing an obvious switch in rates as observed in control.

Surprisingly, we also see a flattening to a single exponential fit with profilin knockdown, suggesting that loss of profilin also slows rapid disassembly (0.105/sec) (Fig 4 C and D, blue line and circle), albeit to a much lesser extent than seen with cofilin knockdown. This effect of profilin on filament disassembly reinforces the interdependency between filament assembly and disassembly, but we don’t have an obvious molecular explanation for this finding. Overall, these results highlight the tight interplay governing filament turnover; where slowing filament assembly slows filament disassembly and reducing filament disassembly by cofilin also slows filament assembly by formin.

### Reduction of components of filament turnover have divergent effects on cortical network integrity and flow

The actomyosin cortex of the one-cell *C.elegans* zygote is composed of filamentous actin that is made up of both long, unbranched actin filaments nucleated by formin/CYK-1 (Severson et al., 2002) and short, branched filaments nucleated by the Arp2/3 complex (Severson et al., 2002; Chan et al., 2019; Xiong et al., 2011). Actin binding proteins such as plastin (PLST-1) organize the cortical F-actin into small bundles that also crosslink F-actin. Myosin II (NMY-2) mini-filaments produce contractile forces on actin filaments that generate both small and large-scale flows in the cortex. Actin architecture of the cortex changes through the cell cycle, coordinated in part through GTPases such as Rho-1 (interphase and cytokinesis) and CDC-42 (mitosis). These changes in actin architecture while maintaining the connectivity of the network generate distinct cortical flow patterns that are required for polarizing the embryo and cytokinesis(Munro et al., 2004; Mayer et al., 2010).

Having quantified the effect of profilin or cofilin knockdown on linear filament assembly and disassembly, we next wanted to evaluate how manipulating turnover affected macroscopic actin dynamics on the network level. When imaging embryos expressing either CYK-1::GFP or actin::GFP we observed an apparent disruption of cortical dynamics and membrane blebbing with knockdown of either cofilin or profilin. To visualize the effect of reducing turnover on cortical actin dynamics, we imaged the actin cortex labeled with utrophin::GFP which binds to, and labels, actin filaments (Video 7) (Michaux et al., 2018; Burkel et al., 2007). Reducing filament disassembly either with strong knockdown of cofilin (UNC-60A), or treatment with 5uM jasplakinolide results in catastrophic tearing of the actin cortex (Video 8). Conversely, slowing formin-mediated filament assembly with strong profilin (PFN-1) knockdown causes contractile instabilities which ultimately also result in tearing of the actin cortex (Video 9), albeit with vastly different dynamics. The complex dynamics of actin cortex collapse make quantification of graded modulation of either filament assembly or disassembly difficult. As a proxy for the evaluation of the effect of turnover on cortex dynamics we focused on non-muscle myosin (NMY-2) flow dynamics at the cortex over the first cell cycle, from interphase to cytokinesis (Fig 5).

**Figure 5.**
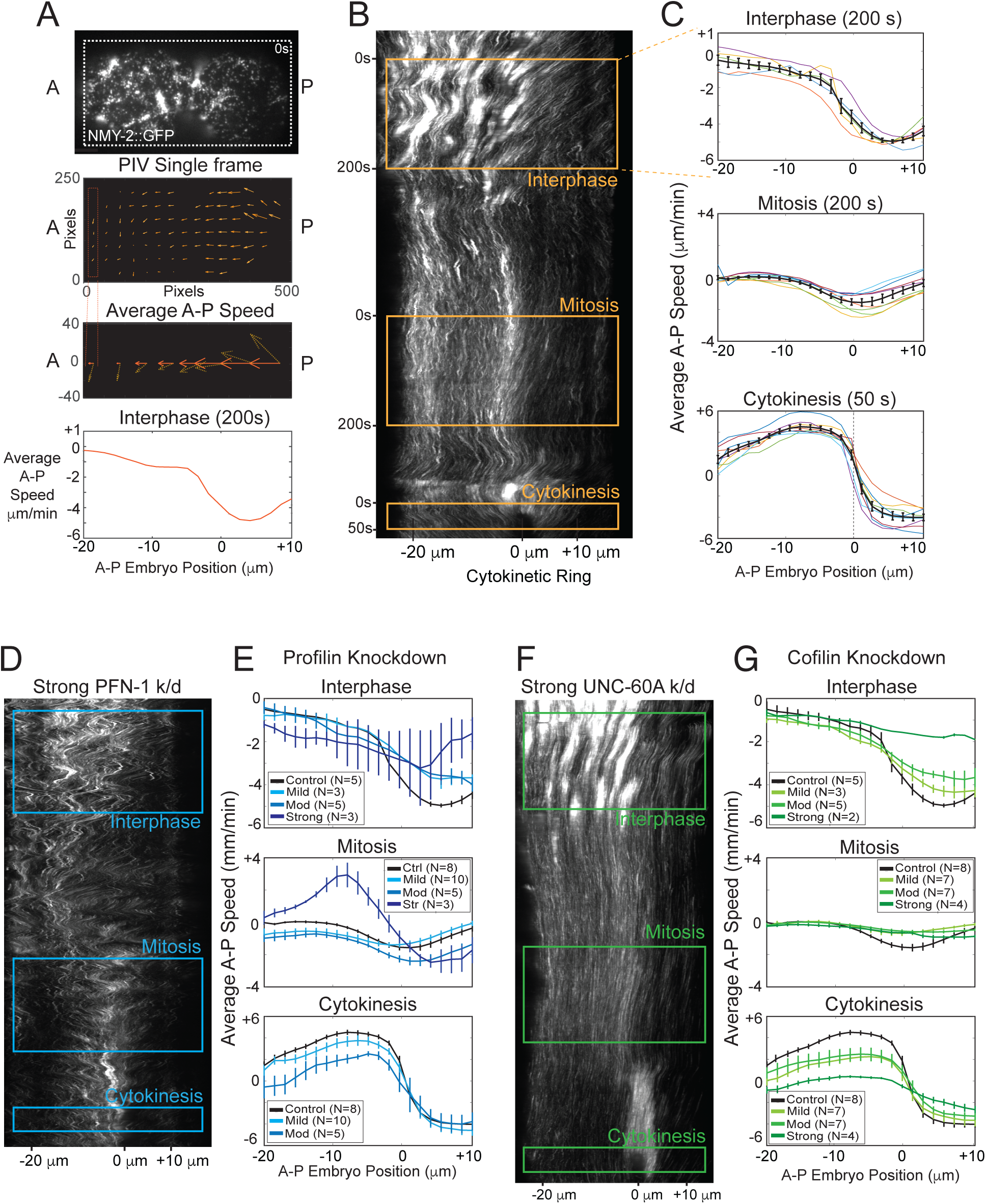
Modifying filament turnover effects myosin flow. **(A)** (Top panel) Surface view of a *C. elegans* zygote in interphase expressing NMY-2::GFP. (Second panel) Quiver plot from PIV analysis of frame in top panel. (Third panel) Averaged axial velocity (*V̅x*) (orange) along the anterior-posterior axis, for the PIV single frame above. (Final panel) Axial velocity versus position along the anterior-posterior axis, averaged over 200 frames during interphase. **(B)** Kymograph of surface view of NMY-2::GFP in the C. elegans zygote over the entire first cell cycle, interphase to cytokinesis. Time is along the y axis. Time windows used for averaged axial velocity plots are indicated by orange boxes on the kymograph. **(C)** Axial velocity versus position profiles along the anterior-posterior axis, for each phase of the cell cycle, interphase (200 seconds), mitosis (200 seconds) and cytokinesis (50 seconds) for 8 control embryos (thin colored lines), mean for all embryos and SEM in black. **(D)** Kymograph of surface view of NMY-2::GFP in the C. elegans zygote for a strong profilin knockdown over the entire first cell cycle, interphase to cytokinesis. **(E)** Axial velocity versus position profiles along the anterior-posterior axis, for each phase of the cell cycle, for graded profilin knockdown (mild (light blue), moderate (blue), and strong (dark blue). Strong knockdown was not included for cytokinesis as most of these embryos do not complete cytokinesis. **(F)** Kymograph of surface view of NMY-2::GFP in the C. elegans zygote for a strong cofilin knockdown over the entire first cell cycle, interphase to cytokinesis. **(G)** Axial velocity versus position profiles along the anterior-posterior axis, for each phase of the cell cycle, for graded cofilin knockdown (mild (light green), moderate (green), and strong (dark green). Error bars are +/- SEM.

To capture cortical flow dynamics, we imaged, via TIRF-M, a transgenically expressed NMY-2::GFP which faithfully recapitulates myosin II dynamics at the cortex (Munro et al., 2004). A representative frame from early in the cell cycle (interphase) displays distinct Rho-A mediated myosin foci at the cortex (Fig 5A, upper panel). Calculated velocity vectors from Particle Image Velocimetry (PIV) of this embryo in interphase is displayed below. These vectors are averaged over the y direction, with the horizontal component plotted over X. To quantify the cortical flow over a representative window in interphase, we selected a 200 second window, with the mean anterior-posterior flow at each position in the embryo plotted in the last panel in A generating a highly reproducible flow pattern characterized by relatively little flow in the anterior of the embryo and strong flows in the posterior of the embryo. Negative values represent posterior to anterior directed flow and positive values represent anterior to posterior directed flow. Using kymography to visualize NMY-2 particle dynamics over the entire cell cycle allows us to see the patterns of cortical flow (Fig 5B). Each time window for interphase (200s), mitosis (200s) or cytokinesis (50s) is outlined in orange, and the corresponding averaged flow profile is shown in C. By selecting a representative time window and averaging over the entire window we can appreciate the robust flow patterns at each stage of the cell cycle. Interphase (Fig 5C, top panel) is characterized by strong anteriorly directed flow that is strongest in the posterior of the embryo. In mitosis (Fig 5C, center panel) cortical flow is significantly reduced so that there is only weak anteriorly directed flow in the posterior of the embryo, and virtually no flow in the anterior of the embryo. Finally, in the 50 second window for cytokinesis, (Fig 5C, bottom panel) there are strong posterior-directed flows in the anterior half of the embryo, and conversely strong anterior-directed flows in the posterior, both converging to zero at the location of the cytokinetic ring.

### Mild and moderate profilin knockdown slows cortical flow, but strong profilin causes increased flow and cortical instabilities

Strong knockdown of profilin causes contractile instabilities, characterized by frequent large-scale rupturing of the actin cortex which can be seen in time-lapse movies (Video 9 (actin) and Video 10 (myosin)), and as rapid anterior-posterior directed movements in kymographs (Fig 5D) at all stages of the cell cycle. In these conditions, the cohesive, uniform cortical flow pattern seen in control (Fig 5C) is lost. Instead, strong reduction of profilin results in erratic flows with significant differences in flow patterns between embryos. We attribute these contractile instabilities to the effect that profilin knockdown has on filament assembly rate (Fig 4B). Shortened linear actin filaments result in the loss of connectivity between the foci of Rho-mediated actin and myosin assembly (Michaux et al., 2018) causing disordered motion of the cortex, which is reflected in the flow profiles quantified by PIV (dark blue lines in Fig 5E). A more nuanced interplay between cortical architecture, filament length and myosin mediated flows can be appreciated by looking at the flow profiles for the graded knockdown of profilin. With mild and moderate profilin knockdown, at times of strong cortical flow, such as in interphase and cytokinesis (Fig 5E, light and medium blue), the pattern of flow is maintained, but the speed of the flow is reduced. This suggests that even small reductions in formin-mediated filament assembly rates, and presumably linear filament lengths, have effects on connectivity and transmission of large-scale flows in the cell. During mitosis, where the relatively mild flows are confined to the posterior of the embryo, the effect of profilin knockdown is more complicated, likely reflecting the change in cortical architecture over the cell cycle. Mild profilin knockdown has relatively little effect on flow, but the moderate knockdown increases flow speeds both in the anterior and posterior of the embryo. This suggests that the cortical architecture during mitosis is organized such that shortened filaments, and reduced connectivity, promotes cortical flow.

### Graded cofilin knockdown slows cortical flow, resulting in cortex collapse

While knockdown of profilin appears to have a complex effect on cortical flow that is dependent on cortex architecture, a simpler picture emerges regarding the role of filament disassembly on cortex dynamics. Mild and moderate knockdown of cofilin both have a relatively mild effect on cortical flows in interphase and cytokinesis, potentially due to the buffering effects of slowed formin assembly rates with cofilin inhibition (Fig 5G). Interestingly, during mitosis when there is less cortical flow, even the mild knockdown of cofilin results in nearly complete loss of flows, which occurs over all knockdown conditions, albeit in a graded fashion, correlating with knockdown strength. This correlates well with the opposing effect that we see with profilin knockdown in mitosis, where shortened, less-connected filaments speed up cortical flow (Fig 5E). Conversely with cofilin knockdown, longer, more crosslinked filaments in the cortex slow cortical flow. With a strong cofilin knockdown (but not as severe as the UTR::GFP knockdown in Video 8) cortical flow and cortex dynamics appear severely reduced at all phases of the cell cycle (Fig 5F and Video 11), suggesting that there is a threshold for the buffering effects of modulating filament length, above which cortex dynamics fail.

Several interesting phenomena can be observed in the video and kymograph of strong knockdown of cofilin, which may give insight into how stabilization of filaments contribute to dissipation of cortical tension and cortex dynamics. Focusing on the myosin dynamics in the anterior of the cortex during mitosis, in the control kymograph (Fig 5B) the myosin particles exhibit small, rapid movements along the anterior-posterior axis, which appear as sinuous paths in the anterior of the embryo. We attribute these patterns to microtears in the cortex that remain locally contained and are not propagated through the cortex and therefore do not generate large scale flows. In contrast, the myosin particles with strong cofilin knockdown have distinctly straight paths (Fig 5F), reflecting the reduced dynamics of the cortex with strong reduction in filament disassembly. With strong knockdown of cofilin, we frequently observe large scale tearing away of the cortex at the posterior end of the embryo throughout all stages of the cell cycle (Video 11), which results in cortex collapse in the most severe cases (Video 8). This large-scale tearing of the cortex contrasts with the pulsatile tearing seen in profilin knockdown, and we interpret the large-scale tearing as due to a failure to dissipate cortical tension (McFadden et al., 2017).

By quantifying myosin flow by PIV, we gained insight into how modulating either filament assembly or disassembly affects cortex dynamics. In general, reduction in filament disassembly slows cortical flows as measured by NMY-2 dynamics. At the strongest knockdown of cofilin, there is a complete collapse and tearing of the cortex. In contrast, strong knockdown of profilin generates chaotic flows, most likely due to loss of cortical connectivity.

### Membrane dynamics reflect the underlying changes in actomyosin cortex behavior

With strong knockdown of either cofilin or profilin, we observed movement of actomyosin cortex components outside the boundary of the ellipsoid embryo, suggesting that the actin cortex is disrupted and membrane blebs may be forming. To examine how F-actin turnover effects membrane dynamics, we performed graded profilin or cofilin knockdowns in strains with lifeact::mCherry to label the actin cortex, and membrane (PLC-d::GFP) (Heppert et al., 2016) to visualize membrane dynamics.

In control embryos, we observe occasional blebbing of the membrane when the actomyosin cortex is undergoing contraction in interphase and cytokinesis (Fig 6A,B and Video 12). These blebs behave as have been previously described in ameboid or cytokinetic animal cells (Charras et al., 2008, 2006; Tinevez et al., 2009) with an initial separation of the membrane from the actin cortex, followed by new assembly of the cortex at the membrane bleb and subsequent flattening of the bleb. One notable difference from these other examples is the speed at which these blebs arise and resolve, which occur within seconds, whereas previously described bleb growth and resolution generally occurs over 10s of seconds (Charras et al., 2008). Unfortunately, due to the speed of the bleb dynamics, it is difficult to collect imaging of both the actin cortex and the membrane over the course of the bleb lifetime due to rapid photobleaching of the actin cortex. Despite this limitation, imaging of the membrane and the cortical actin prior to photobleaching indicates that the described dynamics of bleb formation and resolution occur in the early embryo.

**Figure 6.**
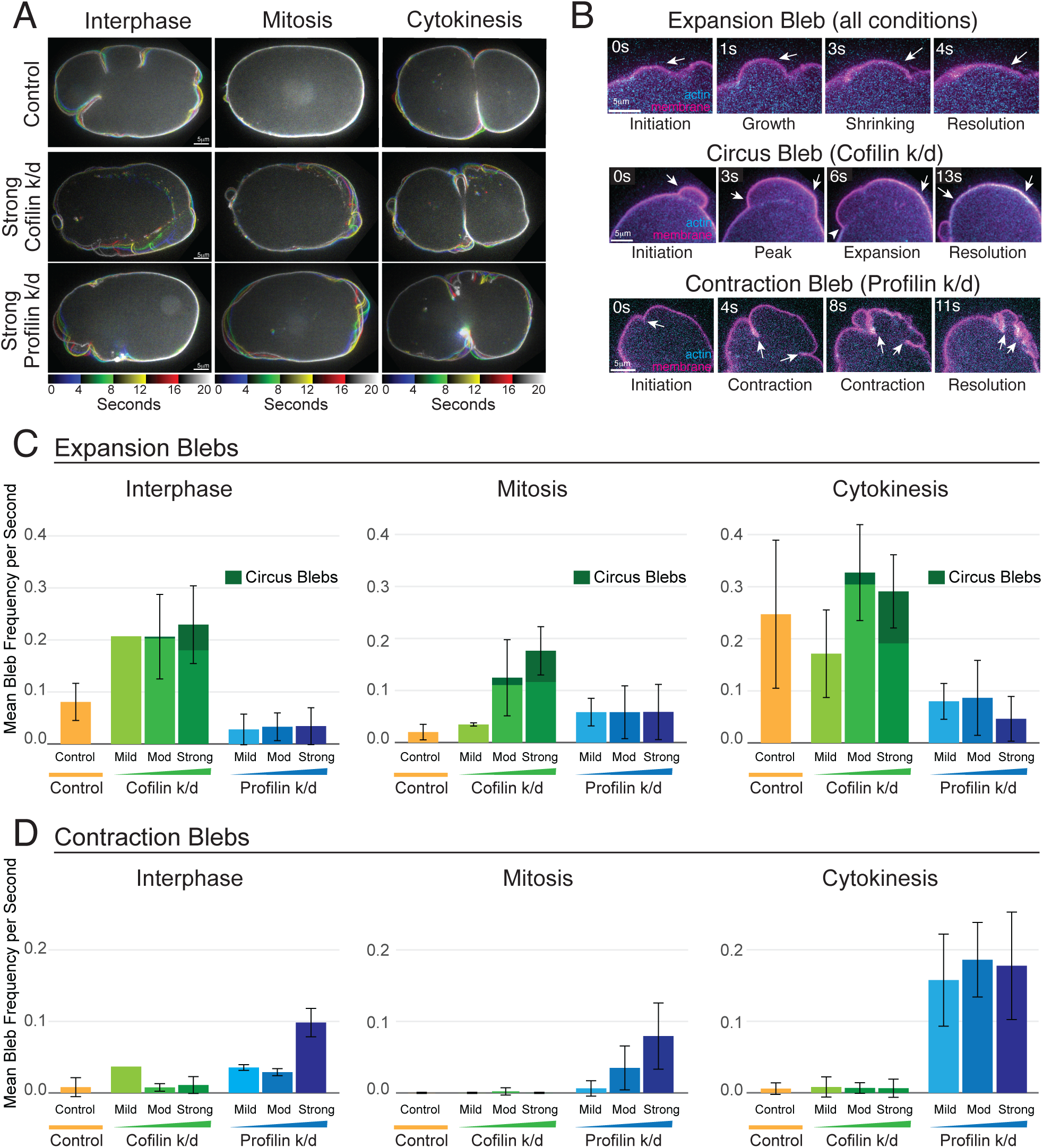
Modifying filament turnover disrupts membrane dynamics. **(A)** Representative mid-section view of a C. elegans zygote in interphase, mitosis and cytokinesis expressing d-PLC::membrane, for control, or strong knockdown of cofilin or strong knockdown of profilin (stills from Video 12). **(B)** All conditions (control, cofilin knockdown and profilin knockdown) exhibit expansion blebs at times of high cortical contractility (interphase and mitosis). Two different types of blebs: circus blebs (seen in cofilin knockdown) and contraction blebs (seen in profilin knockdown) only appear in specific knockdown conditions. **(C)** Quantification of the mean bleb frequency per second for each RNAi knockdown condition, evaluated for each phase of the cell cycle, for expansion blebs. Circus blebs are included as a subset of expansion blebs, indicated in dark green. **(D)** Quantification of contraction blebs for each RNAi knockdown condition, for each phase of the cell cycle.

Both moderate and severe cofilin knockdown result in increased bleb formation in all phases of the cell cycle (Fig 6C), with the strongest cofilin knockdown also resulting in a different type of expansion blebs, termed circus blebs (Fig 6B, middle panels). Circus blebs are characterized by a similar initiation to expansion blebs, but rather than new cortex assembly at the bleb site and resolution, the circus bleb has an additional blebbing of the membrane, further pushing the bleb away from the actomyosin cortex (Charras et al., 2008, 2006). One possibility for this behavior is that in the cofilin knockdown condition, new filament assembly is slowed, so new cortex assembly cannot counteract the internal pressure generating new bleb formation leading to expansion of the bleb rather than resolution. This behavior also highlights the tearing of the cortex that we observe with all actomyosin cortex markers and provides further insight into the mechanics governing this phenomenon.

With knockdown of profilin, we also see an increase in membrane bleb formation, but the dynamics and resolution of these blebs is starkly different from what we observe in either control or reduction in filament disassembly (cofilin knockdown). This is most easily appreciated through imaging of the membrane at the cell surface (Video 12). Using spinning-disc confocal microscopy, a short Z-stack (10μm) at the cell surface is collected and projected. In the control embryo, bleb expansion and resolution can be observed with the membrane returning to a smooth appearance following bleb resolution. In contrast, the cortical tearing that occurs with profilin knockdown (actin imaging Video 9, myosin imaging Video 10) generates an initial stretching of the membrane, then a rapid contraction that generates blebs that get pinched off and do not resolve to a uniform membrane appearance (Figure 6B, Contraction Bleb). We have designated these blebs as contraction blebs to distinguish these bleb dynamics as distinctly different from the traditional bleb dynamics, and reflective of the abnormal actomyosin cortex dynamics caused by knockdown of profilin. We can observe these contraction blebs in cross section (Movie 14 and Fig 6B lower panels) and quantify the frequency of appearance (Fig 6D). While contraction blebs are rarely observed in control and only occasionally observed in cofilin knockdown embryos, contraction blebs are found with both moderate and strong profilin knockdown. As might be predicted, contraction blebs are most frequent at times of high cortical contractility (interphase and cytokinesis), but also appear in mitosis, especially in strong profilin knockdown. By imaging and quantifying membrane dynamics in addition to quantifying myosin dynamics and imaging cortical actin dynamics, we have linked the molecular regulators of actin filament turnover with the large-scale behaviors of the actomyosin cortex and cell membrane.

## Discussion

In this work, we used graded knockdown of proteins involved in actin cortex dynamics to evaluate how modulating the two primary inputs into turnover: rate of assembly and rate of disassembly are tuned to balance remodeling of the actin cortex with maintenance of actomyosin cortex integrity under strain. Using high-resolution TIRF microscopy and single molecule tracking, we were able to quantify, *in vivo*, the effect of reducing turnover on filament assembly and disassembly rates and uncover interdependencies between these inputs. Linking these molecular behaviors with effects on the cellular level, we evaluated both actomyosin cortical flow and membrane dynamics with graded depletion of either profilin or cofilin to determine how slowing either filament assembly or disassembly effects these tightly connected cellular components. We found that disrupting filament turnover effects cortex integrity, cortical flow and membrane dynamics, but that the nature of these disruptions is dependent on if filament assembly or disassembly is disrupted. Despite the inter-dependency between cofilin and profilin, we found that knockdown of either factor had different effects on cortex and membrane dynamics, even if the ultimate result of strong knockdown, failure of cortical integrity and cortex rupture, was the same.

Through careful measurement of filament turnover dynamics, we uncovered both predicted and unexpected results that have the potential to change how we understand actin cortex dynamics in the cell, and as such warrant further discussion. The key findings of this study include: 1) measurement, in vivo, of a delay of about 6 seconds, in rapid actin filament disassembly which would be expected to influence the minimum filament length in the cortex, 2) interdependency between rates of filament assembly and disassembly, where modification of one component of turnover also has effects on the rate of the other factor, and 3) both reduction in linear filament assembly rate and F-actin disassembly rate result in cortex collapse and increase in membrane blebbing. How the cortex tears and collapses in each case is different and careful analysis of these failures provides insight into how these inputs into turnover influence cortical dynamics. These results *in toto* provide a more complete picture of the interplay between cortical turnover and network integrity in vivo and generate new directions for *in vivo*, *in vitro*, and *in silico* investigation.

### Actin filament disassembly has an initial slow rate, followed by rapid dissociation of actin from the cortex

A surprising result from our analysis of actin::GFP lifetime at the cortex was the finding that there are two rates of actin dissociation from the cortex: an initial slow rate that lasts for about 6 seconds (Figure 2G and Figure 3I), followed by a 2-fold faster rate. While we were surprised that our analysis was sensitive enough to observe this *in vivo*, this result fits well with *in vitro* measurements of actin filament disassembly dynamics, where the slow steps of ATP to ADP-Pi conversion and Pi dissociation from the actin polymer precedes rapid actin filament disassembly (Carlier and Pantaloni, 1986; Pollard, 1986; Carlier, 1987; Blanchoin and Pollard, 2002; Jégou et al., 2011). *In vitro*, this slow step of conversion from ATP-actin to ADP-actin can take 300 seconds, much longer than what we observe in vivo. It has long been appreciated that this process of ‘actin aging’ must occur on a faster time scale in the cell, as whole actin network turnover has been measured on the order of seconds (Wang, 1985; Theriot and Mitchison, 1992, 1991; Watanabe and Mitchison, 2002). There is a suite of proteins that are proposed to mediate rapid filament disassembly in the cell, including cofilin, coronin, AIP-1, Srv/CAP, and GMP (Goode et al., 2023). In this study, we focused on cofilin and we found that modulating cofilin concentration via knockdown primarily effects the second, rapid filament disassembly rate but doesn’t seem to significantly change the initial slow disassembly rate, suggesting this initial rate is not dependent on cofilin. Strong knockdown of AIP-1 (AIP) or CAS-1 (Srv/CAP) does not have obvious effects on cortical dynamics in the one-cell embryo (data not shown), but knockdown of POD-1 (coronin) does have a reported phenotype (Rappleye et al., 1999; Xie et al., 2021) and would be an interesting future direction to pursue. In fact, recent work (recent paper) has indicated a prominent role for coronin in facilitating Pi release from filamentous actin.

The observed dynamics of filament disassembly have implications on the architecture of the cortical actin network. With a 6 second delay prior to rapid filament disassembly (Fig 2F), filaments that are assembled by formin, at a rate of 1.2 μm/sec, will have a minimum length of approximately 7.5 μm prior to the start of rapid disassembly, close to the proposed persistence length of actin (10 μm) (Blanchoin et al., 2014). This may indicate that the cortex consists of a sizable proportion of long, linear filaments which would allow for conditions promoting contractility and connectedness required for generation of cohesive cortical flows and cytokinetic ring assembly. This contrasts with a view of the actin cortex consisting of short actin filaments that undergo rapid turnover (Theriot and Mitchison, 1992). It may be that due to the size and polarizing flows of the C. elegans zygote the cortex is specifically tuned to generate longer filaments to perform these cell-specific functions, as such, future work evaluating if there are variations in filament disassembly dynamics in different cell types would be informative. Additionally, using these measured assembly and disassembly parameters in modeling of actin cortex dynamics would allow for more in-depth evaluation how modulating the delay in rapid disassembly effects cortical flow and contractility.

### Inter-dependencies between F-actin assembly and disassembly at the cell cortex

A notable result that became evident through our evaluation of the change in formin-mediated filament assembly rates and filament disassembly rates was the interdependency between these two aspects of filament turnover. As expected, graded reduction in profilin expression though RNAi results in a graded slowing of formin trajectory rates in the embryonic cortex. We would predict that reducing filament turnover by inhibiting cofilin might also slow new filament assembly if there were a limited pool of polymerization competent actin monomers in the cell (Raz-Ben Aroush et al., 2017; Vitriol et al., 2015; Guérin et al., 2025), and our data supports that this occurs *in vivo* (Figure 4B). The fact that reducing disassembly also slows new filament assembly has important implications for how we think about the dynamics of cortex maintenance. If slowing disassembly rates results in concomitant reduction in assembly rates, we might expect that actin filaments in the cortex are maintained at an intermediate length until the buffering effect of slowing filament assembly is exceeded. The cortical flow rates (Fig 5E) and cortical actin dynamics (Video 9) of mild and moderate knockdown of cofilin suggest this buffering effect occurs as the network-level effect of cofilin knockdown appears relatively mild even though there is a significant effect on actin::GFP lifetime at the cortex (Fig 4C).

Perhaps more surprising is the reciprocal relationship between knocking down profilin expression on actin::GFP lifetime at the cortex (Fig4B). As expected, graded knockdown of cofilin significantly increases actin::GFP lifetime at the cortex. Interestingly, we also find that knockdown of profilin increases actin::GFP lifetime at the cortex, albeit not to the degree that we observe with cofilin knockdown. While we do not have an obvious molecular explanation for this result, it does highlight the tight inter-relationship between assembly and disassembly. We note that a synergistic effect of cofilin and profilin on turnover has previously been noted (Didry et al., 1998) from *in vitro* reconstitution assays. One possibility for this effect could be the role that profilin has in ‘recharging’ ADP-actin to ATP-actin, partly by competing with cofilin for ADP-actin (Selden et al., 1999; Mockrin and Korn, 1980; Nishida, 1985). While recent work has highlighted a prominent role for Srv/CAP in this recharging process (Balcer et al., 2003; Kotila et al., 2018), knockdown of Srv/CAP doesn’t have an obvious phenotype in the early *C. elegans* embryo, as such profilin may play a more significant role. In this case, slowed turnover of cofilin-ADP-actin due to decreased profilin may result in reduced availability of cofilin, thereby reducing disassembly. A second possibility is that knockdown of profilin results in increased branched actin assembly, as has been observed in other systems (Burke et al., 2014). Actin::GFP is incorporated into both linear and branched actin networks, and these networks may have different turnover rates. Previous work from our lab supports this possibility, as knockdown of Arp2/3 increases actin::GFP turnover, suggesting that Arp2/3 networks have a slower rate of turnover than formin-generated networks, as has been previously suggested (Fritzsche et al., 2013). Further supporting this possibility, we found a slight increase in the actin::GFP lifetime rate in the anterior of the mitotic cortex, which has increased branched networks as compared to the posterior of the embryo, which has more formin-generated filaments (Supplemental Fig 2). The effect of profilin knockdown on filament lifetime is an area that warrants further study to determine the intricacies of turnover *in vivo*.

### Modifying formin-mediated actin filament assembly, or actin filament disassembly, cause catastrophic loss of cortex integrity

We found that despite the interdependency between filament assembly and disassembly, strong knockdown of either profilin or cofilin had dramatic, but very different effects on cortex integrity, flow, and membrane dynamics. Both perturbations ultimately result in tearing and cortical failure, but the different paths to this result can inform our understanding of factors that shape cortical architecture and dynamics. During interphase and cytokinesis, cortical actomyosin dynamics are dominated by the upstream GTPase Rho, which activates both formin (CYK-1) and myosin (NMY-2), as well as other actin binding proteins (Jaffe and Hall, 2005; Piekny and Glotzer, 2008). This activity generates a cortical network architecture that exhibit large contractile pulses and strong cortical flows (Munro et al., 2004; Michaux et al., 2018; Sailer et al., 2015; Costache et al., 2022; Mayer et al., 2010) (Fig 5C), as well as membrane blebbing. During mitosis, Rho activity decreases and the GTPase CDC42 predominantly shapes network architecture by activating the Arp2/3 complex and myosin II. This results in a change in cortex architecture and myosin activation which results weaker cortical flows in mitosis, and the cell membrane does not exhibit any blebbing. By looking at the effect of turnover on cortical flow and membrane dynamics at these different stages of the cell cycle, we can gain some insight into how the interplay of turnover and actin architecture effect cortical flow and membrane dynamics.

Mild and moderate knockdown of profilin or cofilin both result in slowing of cortical flows in interphase and cytokinesis, but why cortical flow is slowed are different in each case. In the case of profilin knockdown, it appears that shortened linear filaments due to slowed formin-mediated filament assembly decrease the connectivity of the cortex and as a result transmission of contractility and establishment of the large-scale flows is decreased. It is notable that these effects are somewhat moderate until the strongest depletion, which may reflect some amount of buffering of the effect on filament length because of a mild effect on filament disassembly rate.

Cofilin knockdown also slows filament flows, but looking at the videos (Video 11), this appears to be due to the cortex stiffening, and slowing of flows, possibly due to increased connectivity. Despite the buffering effect on filament length that occurs due to slowing of filament assembly rates (Fig 4A), longer filaments might be expected to have increased connections, or the connectivity may be longer lived, resulting in reduced pliancy of the network and reduced elasticity resulting in a reduction in flows. This reduction in fluidity ultimately results in failure to release built-up cortical tension, resulting in tearing of the cortex and the appearance of circus blebs in the strong cofilin knockdown. It is interesting that even with relatively small changes in cortical flow speed observed with moderate cofilin knockdown in interphase and cytokinesis, there is a dramatic increase in the appearance of membrane expansion blebs. We do not think these blebs are due to increased tearing of the cortex, as very little cortical tearing is observed in mild and moderate cofilin knockdown, rather we propose that these blebs are due to either increased cytoplasmic pressure or increased cortical tension due to increased connectivity. Alternatively, the change in cortical dynamics could result in increased breakage of the connections between the cortex and the membrane, resulting in increased frequency of blebbing.

### A special case: The effect of turnover modulation on cortex dynamics in mitosis

The change in cortical architecture and flow dynamics in mitosis provides a window into how these factors are affected by turnover and myosin contraction. In mitosis, both myosin II and the Arp2/3 complex are activated in the anterior of the embryo by CDC-42. This results in very little flow in the anterior of the embryo, likely due to interaction of myosin with the branched actin network (Belmonte et al., 2017; Muresan et al., 2022; Reymann et al., 2012; Alvarado et al., 2013). The posterior of the embryo has more linear filaments, and there is a mild anterior-directed flow in this region (Fig 5B, C, mitosis). This cortical cytoskeleton organization is distinct from Rho-mediated organization and exhibits different effects of slowing either filament assembly or filament disassembly.

In the mitotic actin cell cortex, even mild knockdown of cofilin significantly decreases the anteriorly directed myosin flows. Increased knockdown results in further slowing, until strong knockdown results in tearing of the network in the most posterior of the cell. Additionally, small cortical tears that are observed in the control anterior actin cortex are lost with cofilin knockdown, possibly reflecting a loss of elasticity in the anterior of the cortex, which may be needed for dissipation of cortical tension, as has been suggested by modeling (McCall et al., 2019; McFadden et al., 2017).

In contrast, profilin knockdown in mitosis results in an initial slight slowing of flows, but with increasing reduction of profilin, we observe an increase in flow speeds, that in the strongest knockdown are in both the anterior and posterior of the cell. The initial slowing of flow rates with mild profilin knockdown is likely due to loss of connectivity to transmit flows, as seen in interphase and cytokinesis. The increase in flow with moderate knockdown of profilin may be due to local fast tearing of the cortex dominating in mitosis. In the strongest profilin knockdown, the cortical architecture doesn’t appear to significantly reorganize to the typical mitotic organization, and as such, it is somewhat unclear how this strong phenotype relates to the normal organization of cortical architecture.

During mitosis, membrane blebbing is significantly decreased in control conditions. This may be due to decreased cortical contractility during this phase of the cell cycle. Consistent with this, mild and moderate knockdown of either cofilin or profilin only mildly increases blebbing in mitosis. In contrast, in both conditions strong knockdown significantly increases membrane blebbing, likely due to the observed cortical tearing and failure of cortex integrity.

Specifically evaluating the effects of reducing inputs into actin filament turnover under different cortical architectures and cortex contractile regimes can provide additional insight into how these factors are affected by, and feedback on, filament turnover. Future work specifically modifying actin architecture by inhibiting branched network formation or actin filament bundling will provide additional clarity into network-level effects of turnover on cortical dynamics.

## Conclusion

This work provides a detailed evaluation of how two major inputs into turnover effect cortical actin and membrane dynamics. Building on *in vivo* work from other groups (Naganathan et al., 2018; Mayer et al., 2010; Chugh et al., 2017; Ofer et al., 2011; Ierushalmi et al., 2020; Raz-Ben Aroush et al., 2017) *in vitro* work (Muresan et al., 2022; Murrell and Gardel, 2012; Reymann et al., 2012; Colin et al., 2023), and *in silico* work (McFadden et al., 2017; Yu et al., 2018; Stam et al., 2017; Freedman et al., 2017; Popov et al., 2016; Mak et al., 2016; Belmonte et al., 2017; Jung et al., 2019) this *in vivo* evaluation of actin turnover deepens our understanding of how turnover affects cortex integrity, flow and membrane dynamics. Our careful evaluation provides insights into the feedback and inter-dependency of assembly and disassembly on cortex integrity and cortical flow. Future work examining how actin filament turnover is affected by factors that affect architecture such as bundling and crosslinking actin binding proteins or evaluating dynamics in branched versus linear networks is needed. Using the analyses developed in this paper to allow for careful examination of other inputs into filament length (such as capping protein), or factors that might also affect disassembly rates, such as coronin (Rappleye et al., 1999; Xie et al., 2021; Brieher et al., 2006; Jansen et al., 2015)would provide a more holistic view of how filament length and disassembly affect cortex dynamics.

## Materials and Methods

### C. elegans culture and strains

We cultured the *C. elegans* strains listed in Table 1 under standard conditions (Brenner, 1974) on 60 mm plates containing *E.coli* bacteria (OP50) raised on normal growth medium. Table 1 provides a list of strains used in this study. All strains were cultured at room temperature (20-22°C), except for the actin::GFP strain, which was cultured at 25°C to maintain expression of the actin:GFP transgene. Unless otherwise specified, strains were provided by the Caenorhabditis Genetics Center, which is funded by the National Center for Research Resources.

**Table 1:**
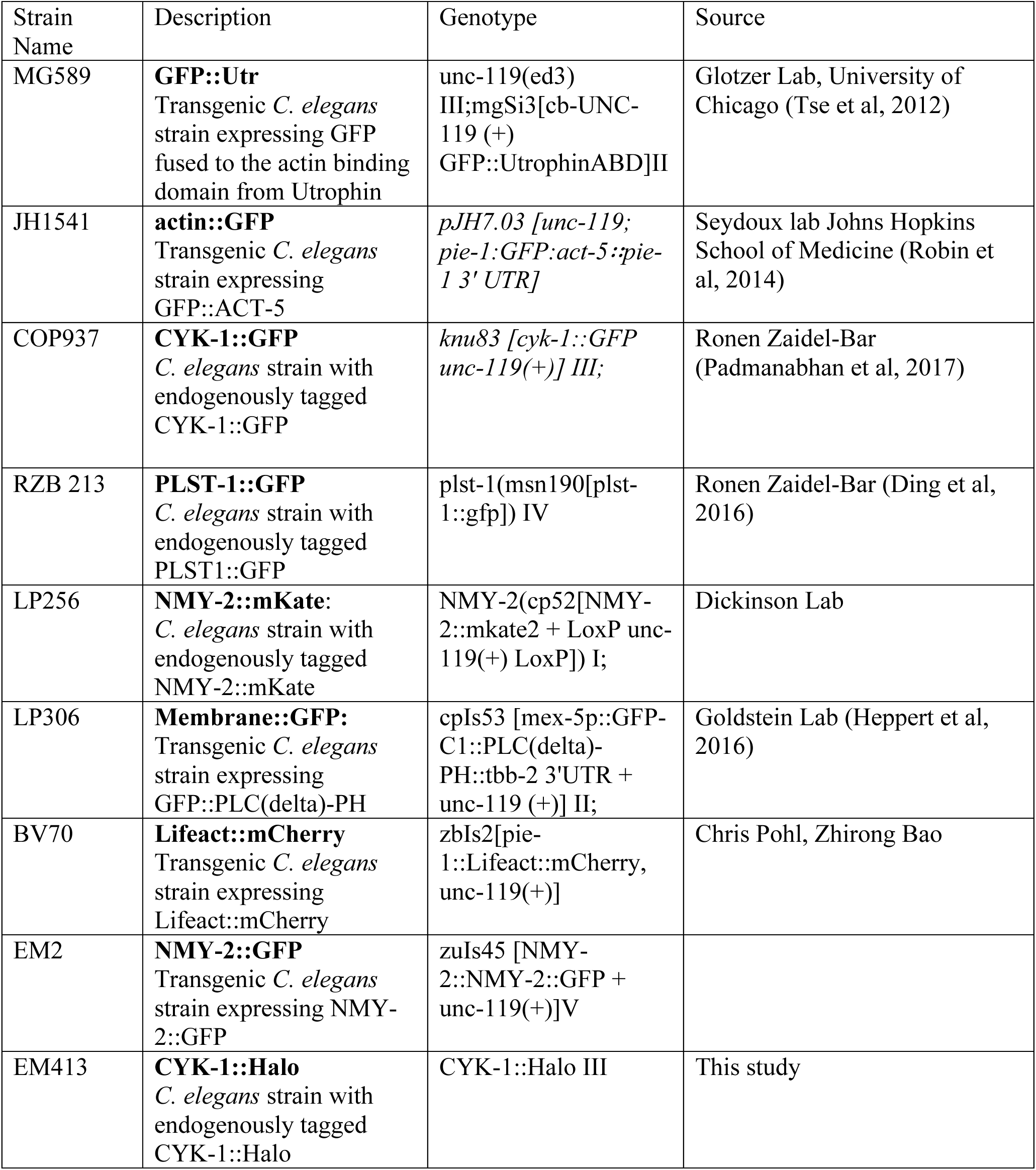
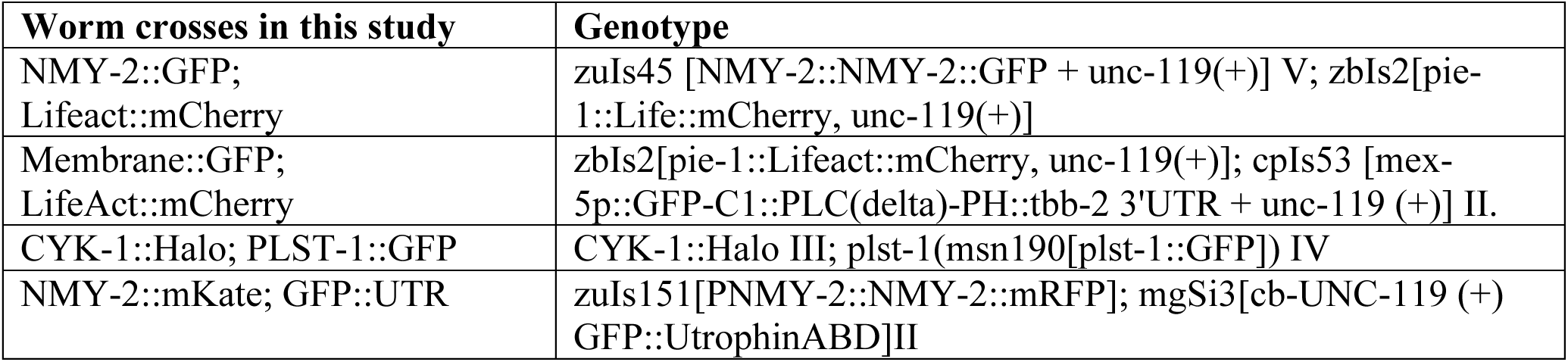
Worm strains used in experiments in this study.

To label the CYK-1::Halo strain with the 647-halo ligand, we cultured L4 worms at 20 degrees in shaking liquid culture for 18 hours with E. coli bacteria supplemented with 50nM conc halo ligand (Chang and Dickinson, 2022).

### RNA interference (RNAi)

We performed RNAi using the feeding method (Timmons et al., 2001). Briefly, we obtained bacteria targeting *UNC-60A, PFN-1* and *NMY-2* from the Ahringer RNAi library (Kamath, 2003). We grew bacteria to log phase in LB with 50μg/ml ampicillin, seeded ∼250μl of culture onto NGM plates supplemented with 50μg/ml ampicillin and 1mM IPTG, incubated plates at room temperature for two days, and then stored them at 4°C for up to five days before use. We transferred L4 larvae to RNAi feeding plates and cultured them at the following temperatures and for the following durations before imaging: 24-30 hours at 20-22 °C for *nmy-2(RNAi)* for strong NMY-2 depletion (Figure 4), 14-26 hours at 20-22 °C for *unc-60A(RNAi)* for mild to strong UNC-60A depletion (Figures 4, 5, 6), 16-26 hours at 20-22 °C for *pfn-1(RNAi)* (Figures 4, 5, 6). For strong depletion of pfn-1 and nmy-2, we first incubated the worms on nmy-2 plates for 6 hours, then moved them to plates with both *pfn-1* and *nmy-2* RNAi (mixed 50:50) for at least 24 hours.

For experiments involving *nmy-2(RNAi)*, we verified strong loss of function by complete failure of first cleavage, and mild loss of function by lack of cortical rotation.

For experiments with a range of *unc-60a* (RNAi), degree of knockdown was correlated with length of time on RNAi plates, and we further confirmed level of knockdown for mild (14-19 hours)- no obvious visual phenotype/difference from control, moderate (19-24 hours)- increase in blebbing, occasional blebbing in mitosis, strong (24-26 hours) increased blebbing and appearance of circus blebs and severe knockdown (>25 hours)- cortex collapse.

For experiments with a range of *pfn-1*(RNAi) degree of knockdown was correlated with length of time on RNAi plates, and we further confirmed level of knockdown for graded mild (16-20 hours)-no obvious visual phenotype, moderate (20-23 hours)- appearance of contractile blebs and contractile instabilities that resolve during mitosis, strong and severe knockdown (>24 hours)-frequent contractile blebs and contractile instabilities throughout the entire cell cycle.

### Live Imaging of *C. Elegans* embryos

We mounted embryos for live imaging in standard egg salts on 2% agarose pads. Embryos were imaged on inverted microscopes for either near-TIRF or spinning disk microscopy (further details below).

### Near-TIRF Microscopy

For timelapse imaging of non-muscle myosin II (NMY-2::GFP), plastin (PLST-1::GFP), or actin (actin::GFP), we used a Nikon Ti2-E inverted microscope equipped with perfect focus and a LunF XL laser combiner with solid state 488 nm, 561 nm, and 640 nm lasers feeding 3 separate TIRF illumination arms. Images were acquired with a 100x 1.49 NA oil-immersion TIRF objective and 1.5X additional magnification onto an Andor IXON-L-897 EM-CCD camera yielding a pixel size of 106 nm. Laser power, illumination direction and angle, and image acquisition were controlled by Nikon Elements software.

For fast near TIRF imaging of CYK-1::GFP, we used an Olympus IX50 inverted microscope equipped with an Olympus OMAC two-color TIRF illumination system, a CRISP autofocus module (Applied Scientific Instrumentation), and a 100x/1.45 NA oil immersion TIRF objective. Laser illumination at 488 nm from a 50-mW solid-state Sapphire laser (Coherent) was delivered by fiber optics to the TIRF illuminator. Images were magnified by 1.6x and collected on an Andor iXon3 897 EMCCD camera, yielding a pixel size of 100 nm. Image acquisition was controlled by Andor IQ software.

For all experiments, we chose a laser illumination angle to maximize the signal-to-noise ratio while maintaining approximately even illumination across the field of view and used the same angle for all observations. Further details of the imaging conditions used for specific quantitative analyses are provided below.

### Spinning Disk Microscopy

For imaging the membrane dynamics in the GFP:PLC-delta; LifeAct::mCherry strain, we used a Nikon ECLIPSE-Ti inverted microscope equipped with Yokogawa CSU-X1 spinning disk and 335 Andor iXon3 897 EMCCD camera, using a 100x/1.49 NA oil immersion objective. For the cell membrane surface images, we acquired Z stacks using x-y motorized stage and fast piezoelectric Z-axis stepper motor with 0.1 μm focus steps. For the mid-section imaging used for quantifying the frequency of bleb appearance, a single z-focal plane at the cell mid-section was obtained every 500ms until the completion of cytokinesis.

### Single-particle detection and tracking

We performed single-particle detection and localization using a MATLAB implementation (http://people.umass.edu/kilfoil/downloads.html) of the Crocker and Grier method (Crocker and Grier, 1996)(Pelletier et al., 2009). Briefly, in each image, the method uses a band pass filter to highlight roughly circular regions below a characteristic size (the feature size), in which the pixel intensity exceeds the background. The regions in which the maximum intensity exceeds a user-defined threshold are identified, and their centroids are determined to sub-pixel resolution as the center of mass of pixel intensity within a pixelated circular mask centered on the original maximum. We used a feature size of 3 and chose thresholds subjectively to optimize detection for different types of particles (actin::GFP or CYK-1::GFP).

Particle-tracking analysis was performed using freely available μTrack software (Jaqaman et al., 2008) (https://github.com/DanuserLab/u-track). μTrack first links particles frame to frame and then links these short segments into longer sequences. Both linking steps use statistical models for particle motion to compute costs for different possible linkage assignments (particle appearance, disappearance, displacement, fusion, and fission) and then identify the assignments that globally minimize these costs. For all analyses reported here, we used a motion model provided with μTrack that represents a mixture of Brownian and directed motion. We allowed the possibility of “gaps” in trajectories due to transient failure to detect particles in individual frames. For each embryo, we overlaid the raw movie with tracked particles to verify tracking accuracy, and we chose parameters for particle detection and tracking (thresholds and length of gaps) that minimize tracking errors. We previously verified the accuracy of these methods for measuring actin filament turnover (Robin et al., 2014). For our analyses of CYK-1 or actin particle movements, all our experiments were performed at particle densities that allowed for identification of individual particles.

### Quantifying CYK-1::GFP particle trajectory speeds at the cortex

We imaged endogenously tagged CYK-1::GFP (Padmanabhan et al, 2017) in streaming mode with 100% laser power and 50 ms exposures and performed particle tracking as previously described(Li and Munro, 2021). We performed our analysis during mitosis, to minimize the effects of cortical flow on trajectory speed, and also allow for quantification of moderate to severe cofilin and profilin knockdown. We selected the subset of trajectories for which log(mean square displacement) vs. log(time) increased with a slope >/= 1.8, consistent with ballistic motion. We confirmed by direct inspection that these trajectories corresponded to rapid directional motion of formin dimers. Because this motion reflects the elongation of single actin filaments, we took the mean speed of these trajectories as an estimate of mean elongation rate. We used (http://github.com/DanuserLab/u-track) developed by Jaqaman and colleagues to track features over time. We integrated particle detection and tracking and all subsequent analyses using custom MATLAB scripts that are available upon request.

### Quantifying actin::GFP lifetime at the cell cortex

To avoid counting false positives – e.g. cytoplasmic molecules that enter and leave the imaging plane without binding, we only considered trajectories with lifetimes >= 2 seconds. We pooled all trajectories collected from multiple embryos at low duty ratio for further analysis. From these pooled data, we constructed single molecule release curves by plotting the number of tracked molecules with lifetime > T as a function of T. We used Matlab’s non-linear least squares fitting function ***nlinfit*** to fit these release curves to two exponential functions, where the interval for the initial rate is smoothly interpolated to the second rate until the best fit is achieved. This generates 3 numbers, the initial observed dissociation rate, the second observed dissociation rate and the switch time. For the streaming, or high duty ratio, data, we constructed single molecule release curves by plotting the number of tracked molecules with lifetime > T as a function of T. We used Matlab’s non-linear least squares fitting function ***nlinfit*** to fit these release curves to a weighted sum of two exponential functions:

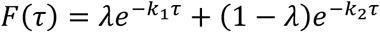

on the interval 0.5 sec < *τ* < ∞, and took the slower of the two rate constants as an estimate of the observed photobleaching rate.

To determine the dissociation rates, which we assume is the sum of rate constant for the dissociation of F-actin-bound from the cortex and the photobleaching rate, which depends on duty ratio:

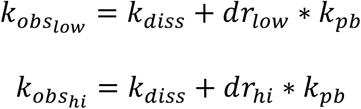

where *k_pb_* is the photobleaching rate during a unit exposure. We solved the above equations simultaneously to estimate the two unknowns (*k_diss_* and *k_pb_*). To compute confidence intervals for these estimates, we used bootstrapping to sample the pooled data 1000 times, computed estimates of *k_diss_* and *k_pb_* from each sample and then took 2 times the standard deviation of the estimates as an estimate of the 95% confidence interval. All scripts used for this analysis are available upon request.

### Plastin streak analysis

We collected a sequence of images from embryos expressing PLST-1::GFP on an Olympus TIRF microscope with 20% laser power and 100 ms exposures. We manually identified individual events in which bright streaks of PLST-1::GFP fluorescence could be observed to grow and then disappear at the actomyosin cortex. In FIJI, we used maximum intensity projections to identify the path of the growing streak; then used the Straighten plugin (https://imagej.net/plugins/straighten) to extract a linear pixel array of width 10 pixels along the path for each frame of the image sequence and converted the resulting image stack into a kymograph for analysis.

To determine plastin spreading speed, a straight line was drawn on the kymograph along the first appearance of the plastin signal. The start and end points of the straight line were determined in Fiji, and velocity calculated from the distance from the end points in x over the distance from the end points in y using pixel size of 0.1 μm and time step of 0.1 seconds.

To determine bundle lifetime, kymographs made as described above for as many streaks as could be identified in each embryo (from 2 to 10 per embryo). The kymograph was manually rotated using the transform feature in Fiji so that the initial appearance of the plastin signal was oriented horizontally. Then the average pixel intensity was calculated for each row in the image, with regions without pixel coverage indicated as NaNs. The change in mean pixel intensity was plotted against time to generate fluorescent intensity profiles for each kymograph (Fig 3F). For each embryo, these intensity profiles were aligned along the initial appearance of plastin signal and then averaged to obtain an averaged intensity profile for each embryo. The profiles for each embryo were then aligned in the same manner and a pooled intensity profile was obtained for all embryos.

### Measuring cortical flow

To measure cortical flows in embryos expressing NMY-2::GFP, we collected data in streaming mode using 20% 488 laser power and 100 msec exposures. Using Fiji ‘grouped Z-project’ with the averaging mode, we averaged 10 frames to generate a movie with 1 second intervals per frame. We measured axial velocity profiles from these data using Particle Image Velocimetry code (mpiv) written in MATLAB (Mori and Chang).

We selected either 200 frames (200 seconds) during interphase and mitosis, or 50 frames (50 seconds) during cytokinesis when flow patterns and speeds show high reproducibility between individual control embryos. To align each velocity profile, we centered each embryo on the location of the cytokinetic ring. In brief we determined the location of the center of the cytokinetic ring and cropped each embryo 10mm posterior and 20mm anterior from the center of the ring. For each frame, we computed a velocity vector field relative to the next frame using the minimum quadratic difference (MQD) algorithm with a sub-window size of 64 pixels and overlap ratio of 0.5 in each direction. We filtered the raw vector field using median filtering and linear interpolation. From the filtered vector field, we computed an axial flow profile by centering each embryo on the location of the cytokinetic ring and extending 10mm towards the posterior and 20 mm towards the anterior. Over this window, we averaged the x (A/P) component of velocity over the y (perpendicular to A/P) direction, then averaged this over time to obtain “steady state” axial flow.

### Quantification of membrane bleb frequency

Because the appearance of blebs is highly correlated with stage of the cell cycle, each video was divided into interphase, mitosis, and cytokinesis for quantification and comparison between knockdown conditions. Blebs that appeared in the imaging plane were counted, with frame of appearance and disappearance noted, as well as location (anterior, mid-line, or posterior). Expansion blebs were defined as an expansion outward of the membrane from the plane of the ellipsoid embryo that exhibited expansion, growth, then resolution with the membrane returning to the shape or location prior to the bleb formation. Circus blebs had the same initial expansion from the membrane, but rather than resolving, additional blebs expanded away from the site of the initial expansion. In the most severe cases, these blebs would move around the circumference of the embryo (exhibiting ‘circus’ motion). Contraction blebs would frequently arise after a stretching of the membrane with a rapid subsequent contraction that resulted in trapping of membrane and incomplete returning of the membrane to its original shape. Any instances where the resolution of the initial bleb resulted in small membrane bubbles that did not fully reintegrate with the membrane were counted as contraction blebs. The bleb frequency was determined by taking the total number of blebs counted in each stage of the cell cycle divided by the number of frames (seconds) in each cell cycle time period.

## Acknowledgements

We thank members of the Kovar and Munro labs for helpful discussions, feedback, and support. In particular, we wish to acknowledge helpful feedback on the manuscript from Kash Baboolall.

This work was supported by National Institutes of Health Grants R35 GM153234 (to D.R.K.), RO1 GM143576 (to E.M.).

